# Complex evolutionary origins of specialized metabolite gene cluster diversity among the plant pathogenic fungi of the *Fusarium graminearum* species complex

**DOI:** 10.1101/639641

**Authors:** Sabina Moser Tralamazza, Liliana Oliveira Rocha, Ursula Oggenfuss, Benedito Corrêa, Daniel Croll

**Affiliations:** Department of Microbiology, Institute of Biomedical Sciences, University of São Paulo, São Paulo, Brazil; Department of Food Science, Food Engineering Faculty, University of Campinas, Av. Monteiro Lobato, 80, Brazil; Laboratory of Evolutionary Genetics, Institute of Biology, University of Neuchatel, Neuchâtel, Switzerland

**Keywords:** head blight, wheat, fungus, pathogen, secondary metabolism

## Abstract

Fungal genomes encode highly organized gene clusters that underlie the production of specialized (or secondary) metabolites. Gene clusters encode key functions to exploit plant hosts or environmental niches. Promiscuous exchange among species and frequent reconfigurations make gene clusters some of the most dynamic elements of fungal genomes. Despite evidence for high diversity in gene cluster content among closely related strains, the microevolutionary processes driving gene cluster gain, loss and neofunctionalization are largely unknown. We analyzed the *Fusarium graminearum* species complex (FGSC) composed of plant pathogens producing potent mycotoxins and causing Fusarium head blight on cereals. We *de novo* assembled genomes of previously uncharacterized FGSC members (two strains of *F. austroamericanum*, *F. cortaderiae* and *F. meridionale*). Our analyses of eight species of the FGSC in addition to 15 other *Fusarium* species identified a pangenome of 54 gene clusters within FGSC. We found that multiple independent losses were a key factor generating extant cluster diversity within the FGSC and the *Fusarium* genus. We identified a modular gene cluster conserved among distantly related fungi, which was likely reconfigured to encode different functions. We also found strong evidence that a rare cluster in FGSC was gained through an ancient horizontal transfer between bacteria and fungi. Chromosomal rearrangements underlying cluster loss were often complex and were likely facilitated by an enrichment in specific transposable elements. Our findings identify important transitory stages in the birth and death process of specialized metabolism gene clusters among very closely related species.

## Introduction

Fungal genomes encode highly organized structures that underlie the capacity to produce specialized (also called secondary) metabolites. The structures are composed of a tightly clustered group of non-homologous genes that in conjunction confer the enzymatic pathway to produce a specific metabolite (Osbourn, 2010). Specialized metabolites (SM) are not essential for the organism’s survival but confer crucial benefits for niche adaptation and host exploitation. Specialized metabolites can promote defense (e.g penicillin), virulence (e.g trichothecenes) or resistance functions (e.g melanin) (Brakhage 1998; Jansen et al. 2006; Nosanchuk and Casadevall 2006). Gene clusters are typically composed of two or more key genes in close physical proximity. The backbone gene encodes for the enzyme defining the class of the produced metabolite and the enzyme is most often a polyketide synthase (PKS), non-ribosomal peptides synthetase (NRPS), terpenes cyclase (TC) or a dimethylallyl tryptophan synthetase (DMATS). Additional genes in clusters encode functions to modify the main metabolite structure (e.g. methyltransferases, acetyltransferases and oxidoreductases), transcription factors involved in the cluster regulation and resistance genes that serve to detoxify the metabolite for the producer (Keller, Turner and Bennet, 2005). The modular nature of gene clusters favored promiscuous exchange among species and frequent reconfiguration of cluster functionalities (Rokas, Wisecaver and Lind, 2018).

The broad availability of fungal genome sequences led to the discovery of a very large number of SM gene clusters (Brakhage, 2013). Yet, how gene clusters are formed or reconfigured to change function over evolutionary time remains poorly understood. The divergent distribution across species (Wisecaver, Slot and Rokas, 2014), frequent rearrangements (Rokas, Wisecaver and Lind, 2018) and high polymorphism within single species (Lind et al. 2017; Wollemberg et al. 2018) complicate the analyses of gene cluster evolution. Most studies analyzed deep evolutionary timescales and focused on the origins and loss of major gene clusters (Wisecaver et al. 2014). Gene clusters often emerged through rearrangement or duplications of native genes (Wong and Wolfe 2005; Slot and Rokas 2010; Wisecaver et al. 2014). The DAL gene cluster involved in the allantoin metabolism is a clear example of this mechanism. The cluster was formed from the duplication of two genes and relocation of four native genes in the yeast *Saccharomyces cerevisae* (Wong and Wolfe 2005). Gene clusters can also arise in species from horizontal gene transfer events (Kaldhi et al. 2008, Khaldi and Wolfe 2011; Campbell et al. 2012; Slot and Rokas 2012). For example, the complete and functional gene cluster underlying the production of the aflatoxin precursor sterigmatocystin was horizontal transferred from *Aspergillus* to the unrelated *Podospora anserine* fungus (Slot and Rokas 2011). Five gene clusters underlying the hallucinogenic psilocybin production were horizontally transmitted among the distantly related fungi *Psilocybe cyanescens, Gymnopilus dilepis* and *Panaeolus cyanescens* (Reynolds et al. 2018). The horizontal transfer was likely favored by the overlapping ecological niche of the involved species.

Despite evidence for high diversity in gene cluster content among closely related strains (Wiemman et al. 2013), the microevolutionary processes driving gene cluster gain, loss and neofunctionalization are largely unknown. Closely related species or species complexes encoding diverse gene clusters are ideal models to reconstruct transitory steps in the evolution of gene clusters. The *Fusarium graminearum* species complex (FGSC) is composed of a series of plant pathogens capable to produce potent mycotoxins and cause the Fusarium head blight disease in cereals. The species complex was originally described as a single species. Based on genealogical concordance phylogenetic species recognition, members of *F. graminearum* were expanded into a species complex (O’Donnel et al. 2004). Currently, the complex includes at least 16 distinct species that vary in aggressiveness, growth rate, and geographical distribution but lack morphological differentiation (Aoki et al. 2012; Ward et al. 2008; Puri and Zhong 2010; Zhang et al. 2012). The genome of *F. graminearum* sensu stricto, the dominant species of the complex, was extensively characterized for the presence of SM gene clusters (Aoki et al. 2012; Wiemman et al. 2013; Proctor et al. 2018; Hoogendoorm et al. 2018). Based on genomics and transcriptomics analyses, Sieber et al. (2014) characterized a large number of clusters with a potential to contribute to virulence and identified likely horizontal gene transfer events.

However, the species complex harbors several other economically relevant species with largely unknown SM production potential (van der Lee et al. 2015). Diversity in metabolic capabilities within the FGSC extends to production of the potent mycotoxin trichothecene. The biosynthesis of some trichothecene variant forms (15-acetyldeoxyvalenol, 3-acetyldeoxynivalenol and nivalenol) are species-specific and associated with pathogenicity (Desjardins et al 2006). Comparative genomics analyses of three species of the complex (*F. graminearum* s.s*, F. asiaticum, F. meridionale*) identified species-specific genes associated with the biosynthesis of metabolites (e.g. PKS40 in *F. asiaticum*) (Walkowiak et al. 2016). Most species were not analyzed at the genome level for SM production potential or lack an assembled genome altogether.

In this study, we aimed to characterize exhaustively the metabolic potential of the FGSC based on comparative genomics analyses and reconstruct the evolutionary processes governing the birth and death process of gene clusters among the recently emerged species. For this, we sequenced and assembled genomes for *F. meridionale*, *F. cortaderiae* and two strains of *F. austroamericanum -* four genomes of the most frequent members of the FGSC found in Brazilian wheat grains, after the well-characterized *F. graminearum* s.s. In total, we analyzed 11 genomes from 8 distinct species within the FGSC. We identified 54 SM gene clusters in the pangenome of the FGSC including two gene clusters not yet known from the complex. The variability in SM gene clusters was generated by multiple independent losses, horizontal gene transfer and chromosomal rearrangements that produced novel gene cluster configurations.

## Material and Methods

### Strains, DNA preparation and sequencing

The fungal strains (*F. meridionale* – Fmer152; *F. cortaderiae* – Fcor153; *F. austroamericanum* – Faus151 and Faus154) were isolated from healthy and freshly harvested wheat grains from three different regions of Brazil, São Paulo State (Fmer152 and Faus151), Parana State (Fcor153) and Rio Grande do Sul State (Faus154) (Tralamazza, et al. 2016). The DNA extraction was performed using a DNAeasy kit (Qiagen, Hilden, Germany) according to the manufacturer’s instructions. DNA quality was analyzed using a NanoDrop2000 (Thermo-Fisher Scientific, USA) and Qubit (Thermo-Fisher Scientific) was used for DNA quantification (minimal DNA concentration of 50 ng/ µL). Nextera Mate Pair Sample Preparation kit (Illumina Inc.) was used for DNA Illumina library preparation. Samples were sequenced using 75 bp reads from paired-end libraries on a NextSeq500 v2 (Illumina Inc.) by the Idengene Inc. (Sao Paulo, Brazil). The software FastQC v. 0.11.7 (Andrews 2010) was used for quality control of the raw sequence reads. To perform phylogenomic analyses, whole genome sequences of *Fusarium* species and *Trichoderma reesei* (as an outgroup) were retrieved from public databases (see Supplementary Table S1 for accession numbers).

### Genome assembly

D*e novo* genome assembly was performed for the four newly sequenced genomes of the FGSC (*F. meridionale* – Fmer152; *F. cortaderiae* – Fcor153; *F. austroamericanum* – Faus151 and Faus154) and for the publicly available 150 bp paired-end raw sequence data for *F. boothi*, *F. gerlachii* and *F. louisianense* (Supplementary Table S1). We used the software Spades v.3.12.0 (Bankevich et al. 2012) to assemble Illumina short read data to scaffolds using the “careful” option to reduce mismatches. We selected the k-mer series “21,33,45,67” for *F. meridionale, F. cortaderiae* and *F. austroamericanum* sequences, and “21,33,55,77,99,127” for *F. boothi*, *F. gerlachii* and *F. louisianense*. The maximum k-mer values were adjusted according to available read length. For all other genomes included in the study (including *F. asiaticum* and *F. graminearum* s.s), assembled scaffolds were retrieved from NCBI or Ensembl database (Supplementary Table S1). The quality of draft genome assemblies was assessed using QUAST v.4.6.3 (Gurevich et al. 2013). BUSCO v.3 (Waterhouse et al. 2017) was used to assess the completeness of core fungal orthologs based on a fungal BUSCO database.

### Gene prediction and annotation

Genes were predicted using Augustus v.2.5.5 (Stanke and Morgenstern 2005). We used the pre-trained gene prediction database for the *F. graminearum* s.s genome as provided by the Augustus distribution for all annotations and used default parameters otherwise. Predicted proteomes were annotated using InterProScan v.5.19 (Joones et al. 2014) identifying conserved protein domains and gene ontology. Secreted proteins were defined according to the absence of transmembrane domains and the presence of a signal peptide based on Phobius v.1.01 (Kall et al. 2004), SignalP v.4.1 (Petersen et al. 2011) and TMHMM v.2.0 (Krog et al. 2001) concordant results. We identified the predicted secretome with a machine learning approach implemented in EffectorP v2.0 (Sperschneider et al. 2018).

### Genome alignment and phylogenomic analyses

For the phylogenomic analyses, we used OrthoMCL (Li et al. 2003) to identify single copy orthologs conserved among all strains. High accuracy alignment of orthologous sequences was performed using MAFFT v.7.3 (Katoh et al. 2017) with parameters --maxiterate 1000 --localpair. To construct a maximum-likelihood phylogenetic tree for each alignment, we used RAxML v.8.2.12 (Stamatakis 2014) with parameters -m PROTGAMMAAUTO and bootstrap of 100 replicates). The whole-genome phylogeny tree was constructed using Astral III v.5.1.1 (Zhang et al. 2017) which uses the multi-species coalescent model and estimates a species tree given a set of unrooted gene trees. We used Figtree v.1.4.0 for visualization of phylogenetic trees (Rambaut 2012).

### Specialized metabolite gene cluster prediction

To retrieve specialized metabolite (SM) gene clusters from genome assemblies, we performed analyses using antiSMASH v.3.0 (Blin et al. 2017) and matched predicted gene clusters with functional predictions based on InterProScan v. 5.29-68 (Jones et al. 2014). For the *F. graminearum* reference genome (FgramR), we retrieved SM gene clusters identified in a previous study, which used evidence from multiple prediction tools and incorporated expression data (Sieber et al. 2014). We selected only clusters with a defined class/function, identified backbone gene and annotated cluster size. We made an exception for cluster SM45, which was predicted by antiSMASH but not characterized by Sieber et al. (2014) likely due to discrepancies in gene annotation.

### Pangenome SM gene cluster map and synteny analysis

We constructed a pangenome of SM gene clusters in the FGSC by mapping the backbone genes of each distinct cluster against all other genomes. BLAST+ v.2.8 (Camacho et al. 2009) local alignment search (blastp with default parameters) was performed and matches with the highest bitscores were retrieved. For each unique cluster in FGSC, we selected the backbone gene of a specific genome as a reference for presence/absence analyses within the complex. We used FgramR backbone sequences for the majority of the clusters (clusters SM1-SM45), for SM46 we used FasirR2, for SM47-SM52 FasiR, for SM53 we used Fcor153 and for SM54 we used Faus154 (Supplementary Table S3). We considered a gene cluster as present if the blastp identity of the backbone gene was above 90% (threshold for FGSC members). For strains outside of the FGSC (*i.e.* all other *Fusarium* species), we used a cut-off of 70%. Heatmaps were drawn using the R package ggplot2 (Wickham 2016) and syntenic regions of the gene clusters were drawn using the R package genoplotR (Guy et al. 2010). For SMGC with taxonomical distribution mismatching the species phylogeny, we performed additional phylogenetic analyses. For this, we queried each encoded protein of a cluster in the NCBI protein database (see Supplementary Table S2 for accession numbers). We reconstructed the most likely evolutionary history of a gene cluster using the maximum likelihood method based on the JTT matrix-based amino acid substitution model (Jones et al. 1992). We performed 1000 bootstrap replicates and performed all analyses using the software MEGA v.7.0.26 (Kumar et al. 2016).

### Repetitive elements annotation

We performed *de novo* repetitive element identification of the complete genome of *F. graminearum* (FgramR) using RepeatModeler 1.0.11 (Smit and Hubley 2008). We identified conserved domains of the coding region of the transposable elements using blastx and the non-redundant NCBI protein database. One predicted transposable element family was excluded due to the high sequence similarity to a major facilitator superfamily gene and low copy number (*n* = 2), which strongly suggests that a duplicated gene was misidentified as a transposable element. We then annotated the repetitive elements with RepeatMasker v.4.0.7 (Smith et al. 2015). One predicted transposable element family (element 4-family1242) showed extreme length polymorphism between the individual insertions and no clearly identifiable conservation among all copies. The consensus sequence of family1242 also contained several large poly-A islands, tandem repeats and palindromes. Using blastn, we mapped the sequences of all predicted insertions against the consensus sequence and identified five distinct regions with low sequence similarity between them. We created new consensus sequences for each of these five regions based on the genomes of *F. graminearum* and *F. austroamericanum* (Faus154) (Morgulis et al. 2008; Zhang et al. 2000). We filtered all retrieved sequences for identity >80% and >80% alignment length. We added flanking sequences of 3000 bp and visually inspected all retrieved hits with Dotter v.3.1 (Sonnhammer and Durbin 1995). Then, we performed a multiple sequence alignment using Clustalw (Altschul et al. 1997; Higgins and Sharp 1988) to create new consensus sequences. Finally, we replaced the erroneous element 4-family 1242 with the five identified sub-regions. We used the modified repeat element library jointly with the Dfam and Repbase database to annotate all genomes using RepeatMasker (Smit et al. 2008). Transposable element locations in the genome were visualized with the R package genoPlotR v0.8.9 (Guy et al. 2011). We performed transposable element density analyses of the genomes in 10 kb windows using bedtools v.2.27 (Quinlan and Hall 2010).

## Results

### Genomic sampling of the *Fusarium graminearum* species complex

We analyzed genomes of 11 strains of 8 different species of the FGSC in order to resolve species relationships and detect divergence in their specialized metabolism. We performed the first *de novo* assembly and genome annotation for two strains of *F. austroamericanum* (Faus151 and Faus154), a strain of *F. cortaderiae* (Fcor153) and a strain of *F. meridionale* (Fmer152). We included 15 other species of the *Fusarium* genus including the *Fusarium fujikuroi* species complex (FFSC) and the *Fusarium sambucinum* species complex (FSAMSC) to distinguish between gene gains and losses. We first assessed the genome assembly quality within FGSC (Supplementary Table S1). N50 values of the newly sequenced genomes ranged from 220-442 kb. The N50 of previously sequenced genomes of the FGSC ranged from 149-9395 kb including the fully finished assembly of the reference genome *F. graminearum* PH-1 (FgramR). By analyzing the completeness of all assemblies, we found the percentage of recovered BUSCO orthologues to be above 99.3% for all FGSC members. The genome sizes within the FGSC ranged from 35.02 – 38.0 Mbp. All genomes shared a similar GC content (47.84 – 48.39%) and number of predicted genes (11’484-11’985) excluding the reference genome. The *F. graminearum* reference genome showed a higher number of predicted genes (14’145) most likely due to the completeness of the assembly and different gene annotation procedures. The percentage of repetitive elements in the genome varied from 0.47 – 4.85% among members of the *Fusarium* genus with a range of 0.97 – 1.99% within the FGSC. Genomes of strains falling outside of the FGSC showed N50 values and a BUSCO recovery of 31–9395 kb and 93– 100%, respectively.

### Phylogenomic reconstruction

We analyzed the phylogenetic relationships of eight distinct species within the FGSC and 15 additional members of *Fusarium*. We included *Trichoderma reesei* as an outgroup species. Using OrthoMCL, we identified 4191 single-copy orthologs conserved in all strains and used these to generate a maximum likelihood phylogenomic tree (Figure 1). The three species complexes included in our analyses (FFSC, FSAMSC and FGSC) were clearly differentiated with high bootstrap support (100%). All FGSC members clustered as a monophyletic group and *F. culmorum* was the closest species outside of the complex. The cluster of *F. graminearum, F. boothi, F. gerlachii* and *F. louisianense*, as well *F. cortaderiae, F. austroamericanum* and *F. meridionale* each formed well-supported clades. The FGSC species clustered together consistent with previous multi-locus phylogenetic studies based on 11 combined genes (Aoki et al. 2012) apart from *F. asiaticum* clade that was found separated from the clade of *F. graminearum, F. boothi, F. gerlachii* and *F. louisianense*. The tree clearly resolves the FSAMSC as a monophyletic group, which includes *F. culmorum, F. pseudograminearum, F. langsethiae, F. poae* and *F. sambucinum*, together with all members of the FGSC. The members of the FFSC (*F. fujikuroi, F. verticillioides, F. bulbicola, F. proliferatum* and *F. mangiferae*) also formed a monophyletic group.

**Figure 1.**
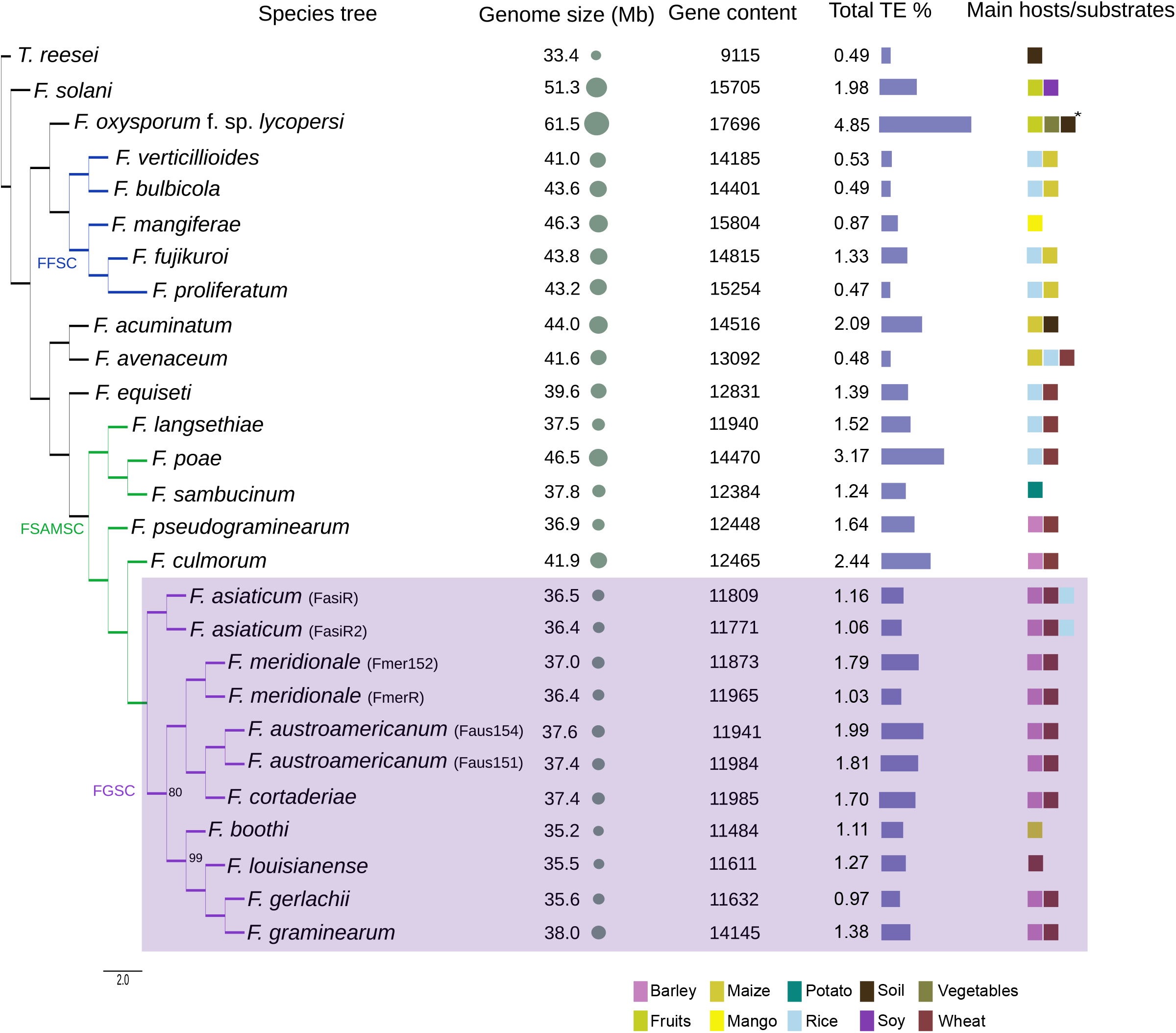
Phylogenomic tree of the *Fusarium graminearum* species complex (FGSC) and other *Fusarium* strains, inferred from a coalescence-based analysis of 4192 single-copy orthologues and bootstrap of 100 replicates. *T. reseei* was used as an outgroup. Tree nodes without values have a bootstrap of 100%. Substrate/host information was retrieved from Aoki et al. (2012). * *F. oxysporum* lineages are usually host specific. FFSC: *Fusarium fujikuroi* species complex. FSAMSC: *Fusarium sambucinum* species complex.

### Specialized metabolite gene clusters diversity in the FGSC

We analyzed all genome assemblies for evidence of SM gene clusters based on physical clustering and homology-based inference of encoded functions. Out of 54 SM gene cluster within the FGSC, seven were absent from the *F. graminearum* reference (Figure 2). The class of NRPS was the most frequent SM gene cluster category (*n* = 19), followed by PKS (*n* = 13) and TPS (*n* = 11). We also found several cases of hybrid clusters, containing more than one class of backbone gene (Figure 2). We found substantial variation in the presence or absence of SM gene clusters within the FGSC and among *Fusarium* species in general. We classified gene clusters into three distinct categories based on the phylogenetic conservation of the backbone gene in FGSC (Figure 2). Out of the 54 clusters, 43 SM gene clusters were common to all FGSC members (category 1; Figure 2). The SM gene clusters shared within the species complex were usually also found in the heterothallic species *F. culmorum* (86.4% of all clusters) and in *F. pseudograminearum* (79.7% of all clusters), the most closely related species outside of the FGSC (Figure 1). The gene cluster responsible for the production of the metabolite gramillin was shared among all FGSC species and *F. culmorum* (Figure 2). We found five SM gene clusters (SM22, SM43, SM45 and SM48) that were not shared by all FGSC members but present in more than 20% of the strains (category 2; Figure 2). Six SM gene clusters (SM46, SM50, SM51, SM52, SM53 and SM54) were rare within the FGSC or even unique to one analyzed genome (category 3; Figure 2). We also found 13 highly conserved SM gene clusters among members of the *Fusarium* genus with 24 of the 26 analyzed genomes encoding the backbone gene (>70% amino acid identity; Supplementary Table S3). An example of such a conserved cluster is SM8 underlying the production of the siderophore triacetylfusarine, which facilitates iron acquisition both in fungi and bacteria (Charlang et al. 1981).

**Figure 2.**
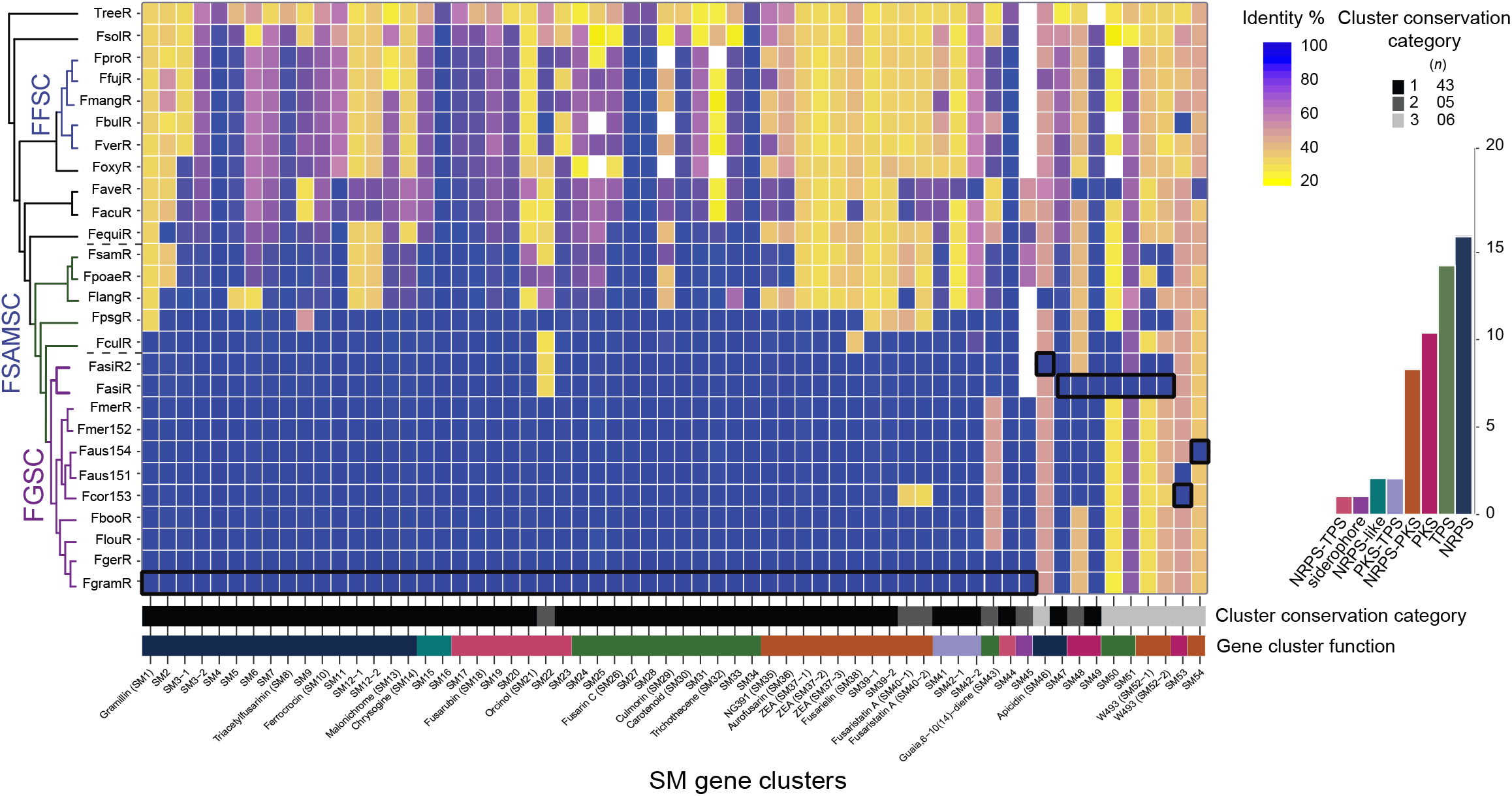
Secondary metabolite gene cluster pangenome of the *Fusarium graminearum* species complex (FGSC) based on evidence for backbone genes. Squares with black lines in the heatmap correspond to genomes used for comparative genomic analyses: FgramR (SM1-SM45), FasiR2 (SM46), FasiR (SM47-SM52), Fcor153 (SM53) and Faus154 (SM54). The bar chart is identifying frequencies clusters types. Colored bars below the heatmap correspond to the cluster type. Black/grey bars correspond to the category conservation cluster table. PKS – polyketide synthase; NRPS – nonribosomal peptide synthetase; TPS – terpene synthase. FFSC: *Fusarium fujikuroi* species complex. FSAMSC: *Fusarium sambucinum* species complex

### Multiple gene cluster rearrangements and losses within the FGSC

We analyzed the mechanisms underlying gene cluster presence-absence polymorphism within the FGSC (category 2 and 3; Figure 2). These clusters were encoding the machinery for the production of both known and uncharacterized metabolites. We considered a gene cluster to be lost if at least the backbone gene was missing or suffered pseudogenization. Both, SM45, underlying siderophore production, and SM33, a PKS cluster, were shared among all FGSC members except *F. asiaticum* (FasiR). The cluster of fusaristatin A (SM40), a metabolite with antibiotic activities and expression associated with infection in wheat (Sieber et al. 2014) was another example of cluster loss in a single species, *F. cortaderiae* (Fcor153). We found that the cluster encoding for the production of the metabolite guaia,6-10(14)-diene (SM43) is conserved in different species within FGSC but the cluster suffered independent losses in *Fusarium*. The TPS class gene cluster identified in *F. fujikuroi* (Burkhardt et al. 2016) was shared among different species complexes (FFSC and FSAMSC; Figure 3). In the FFSC, the species *F. fujikuroi, F. proliferatum, F. bulbicola* and *F. mangiferae* share the cluster. In the FSAMSC, the parent complex that includes also FGSC, the guaia,6-10(14)-diene cluster was found to be rearranged compared to the cluster variant found in the FFSC. Gene cluster synteny analyses among strains within the FGSC showed that several members (*F. cortaderiae, F. austroamericanum, F. meridionale* and *F. louisianense*) lost two segments of the cluster. The gene cluster variant with partial deletions retained only the gene encoding for the biosynthesis of pyoverdine and the genes flanking the cluster (Figure 3). To retrace the evolutionary origins of the guaia, 6-10(14)-diene cluster, we performed a phylogenetic analysis of each gene within the cluster. The backbone gene encoding for the terpene synthase and the pyoverdine biosynthesis genes show congruent phylogenetic relationships. However, the gene phylogenies showed discrepancies compared to the species tree (Supplementary Figure S1). Both gene trees showed that orthologs found within the FGSC grouped with species outside of the complex. *F. graminearum* and *F. gerlachii* formed a subclade with the sister species *F. culmorum* as did *F. asiaticum* with the FSAMSC species *F. pseudograminearum*.

**Figure 3.**
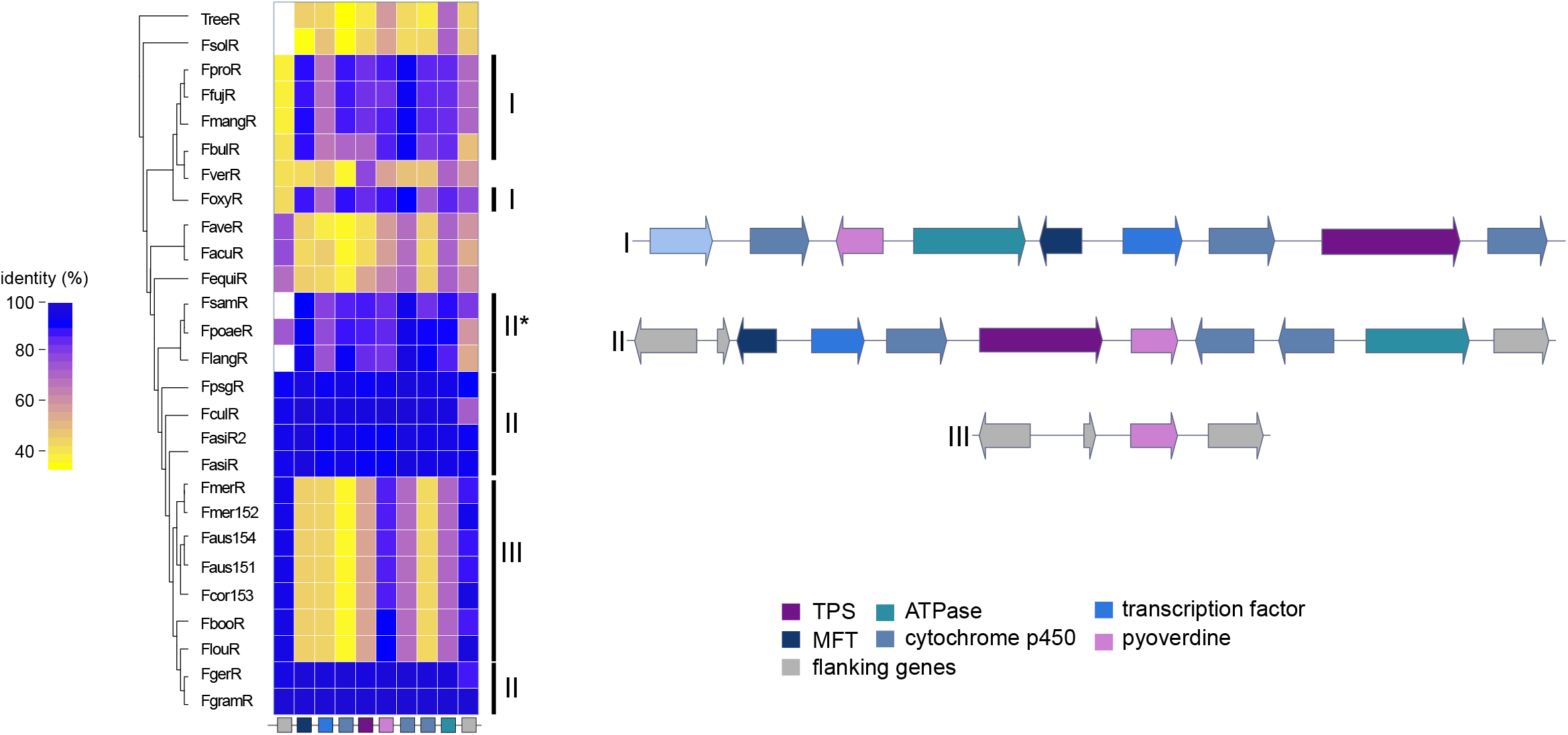
Synteny plot of the SM46 (guaia-6,10-diene) gene cluster and heatmap of protein identity based on the *Fusarium graminearum* FgramR reference genome. Rectangles below the heatmap correspond to the genes shown in the synteny plot. Arrows of identical color correspond to homologous genes and identify the predicted protein function. TPS: terpene synthase; MFT: major facilitator superfamily transporter.

We found the cluster underlying the apicidin metabolite production (SM46) present within the FGSC (Figure 4). The cluster was first discovered in *F. incarnatum* (former *F. semitectum*; Jin et al. 2010) and was found to underlie the production of metabolites with antiparasitic proprieties (Darkyn-Ratway et al. 1996). Our analysis showed that the cluster suffered multiple independent losses across the *Fusarium* genus including a near complete loss within the FGSC, except in the strain of *F. asiaticum* (FasiR2), which shares a complete and syntenic cluster with the distantly related species *F. incarnatum* and *F. langsethiae*. Surprisingly, the *F. asiaticum* strain FasiR maintained only a pseudogenized NRPS backbone gene and the flanking genes on one end of the cluster. *F. fujikuroi* is missing *aps10* encoding a ketoreductase and is known to produce a similar metabolite called apicidin F (Niehaus et al. 2014). We performed a phylogenetic analysis of the genes *aps1* encoding an NRPS, *aps5* encoding a transcription factor, *aps10* and *aps11* encoding a fatty acid synthase to investigate a scenario of horizontal gene transfer. Both the individual gene trees and a concatenated tree (with *aps1*, *aps5* and *aps11*) showed that the genes follow the species tree phylogeny except for *F. avenaceum* (Figure 4). The phylogeny of *aps10* included a homologous gene of *F. acuminatum*, which together with *F. avenaceum*, is part of the *Fusarium tricinctum* species complex. The phylogeny of *aps10* diverged from the species tree. An analysis of gene cluster synteny showed that the *F. avenaceum* gene cluster is missing the gene *aps9* and underwent a drastic gene order rearrangement compared to the other species. The rearrangement and divergency may be the consequence of a partial gene cluster duplication and may have led to a neofunctionalization of the gene cluster in *F. avenaceaum*. The sequence rearrangement in the apicidin gene cluster and the discontinuous taxonomic distribution is suggestive of a horizontal gene transfer event from *F. langsethiae* to *F. asiaticum.* However, multiple independent losses across the *Fusarium* genus combined with a possible advantage to maintain the cluster in the *F. asiaticum* strain FasiR2 could explain the observed patterns as well (Figure 4).

**Figure 4.**
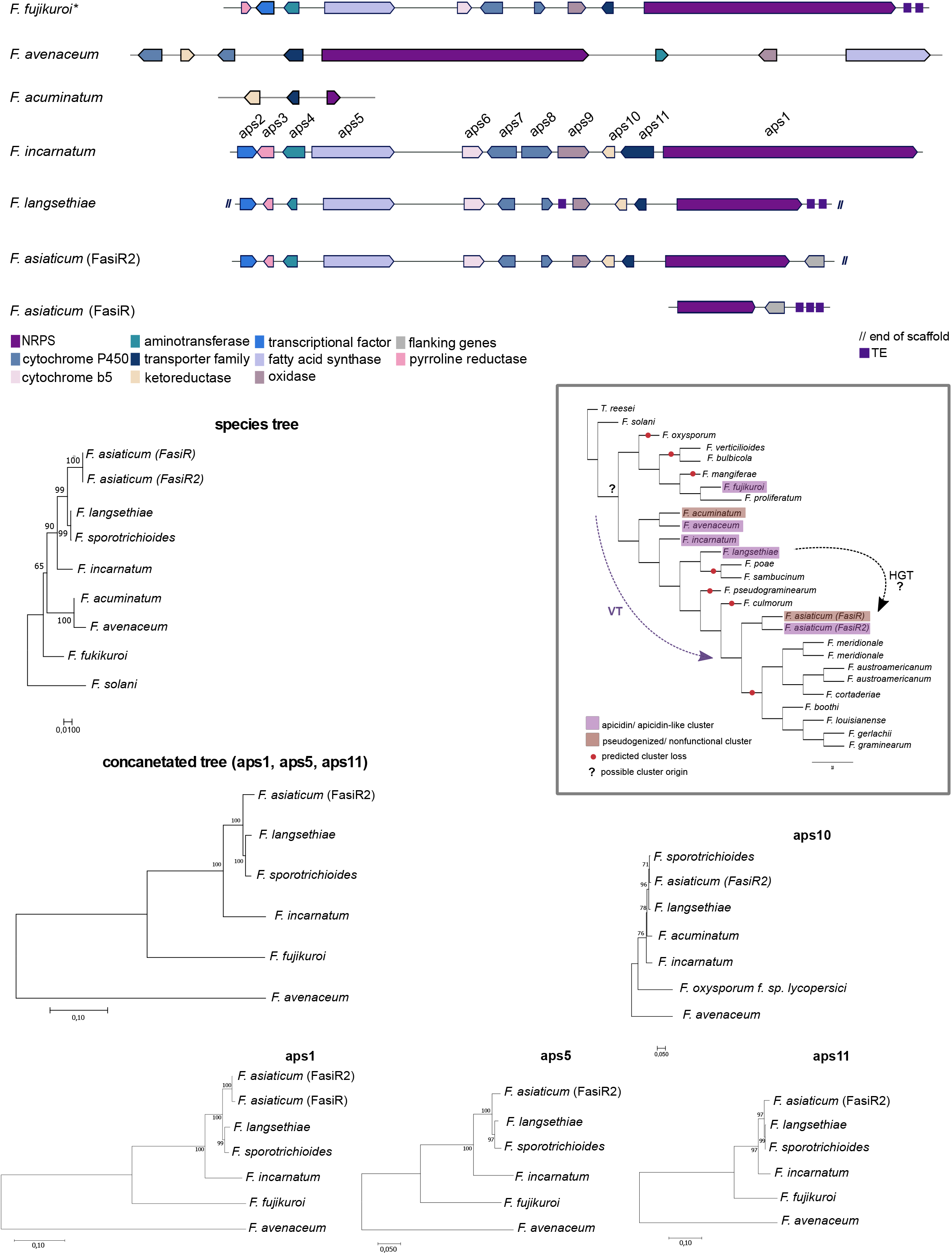
Synteny plot of the SM46 apicidin metabolite gene cluster. Arrows of identical color correspond to homologous genes and identify the predicted protein function. * Fusarium fujikuroi is an apicidin-F producer. Phylogenetic trees were constructed using maximum likelihood and the JTT matrix-based amino acid model with 1000 bootstrap replicates. The species tree was based on the concatenated analysis of the EF-1a, RBP1 and RPB2 genes. *Fusarium solani* was used as the outgroup. Grey boxes indicate the presence, independent loss and possible origin of the apicidin cluster. HGT: horizontal gene transfer. VT: vertical transmission.

### A secondary gene cluster is linked to multiple horizontal gene transfers events

We found evidence for a horizontal transfer of six genes among fungi and a single bacterial transfer event in the formation of the SM54 gene cluster. The rare cluster (category 3), with a predicted size of 11 genes, was found in the FGSC strain *F. austroamericanum* (Faus154). Across *Fusarium* species, six genes of the cluster are shared with *F. avenaceum* (Figure 5). Of the six genes, the backbone gene encoding the PKS, a cytochrome P450 and a methyltransferase gene share homology with the genes *fdsS, fdsH* and *fdsD*, respectively, constituting the Fusaridione A cluster in *F. heterosporum*. A homology search of the genes shared between *F. austroamericanum* and *F. avenaceum* showed *F. avenaceum* to be the only hit with a high percentage of identity (>80%) to the analyzed genes (Supplementary Table S4). The phylogenetic analyses of the six genes, consistently grouped *F. austroamericanum* with *F. avenaceum.* This clustering was conserved if the tree included also orthologs found in *F. heterosporum*, which is a species more closely related to *F. avenaceum* than *F. austroamericanum* (Figure 5). The phylogenetic distribution of the gene cluster and high homology strongly suggest that at least a segment of the cluster was horizontally transferred from the *F. avenaceum* lineage to *F. austroamericanum* to create the SM54 gene cluster.

**Figure 5.**
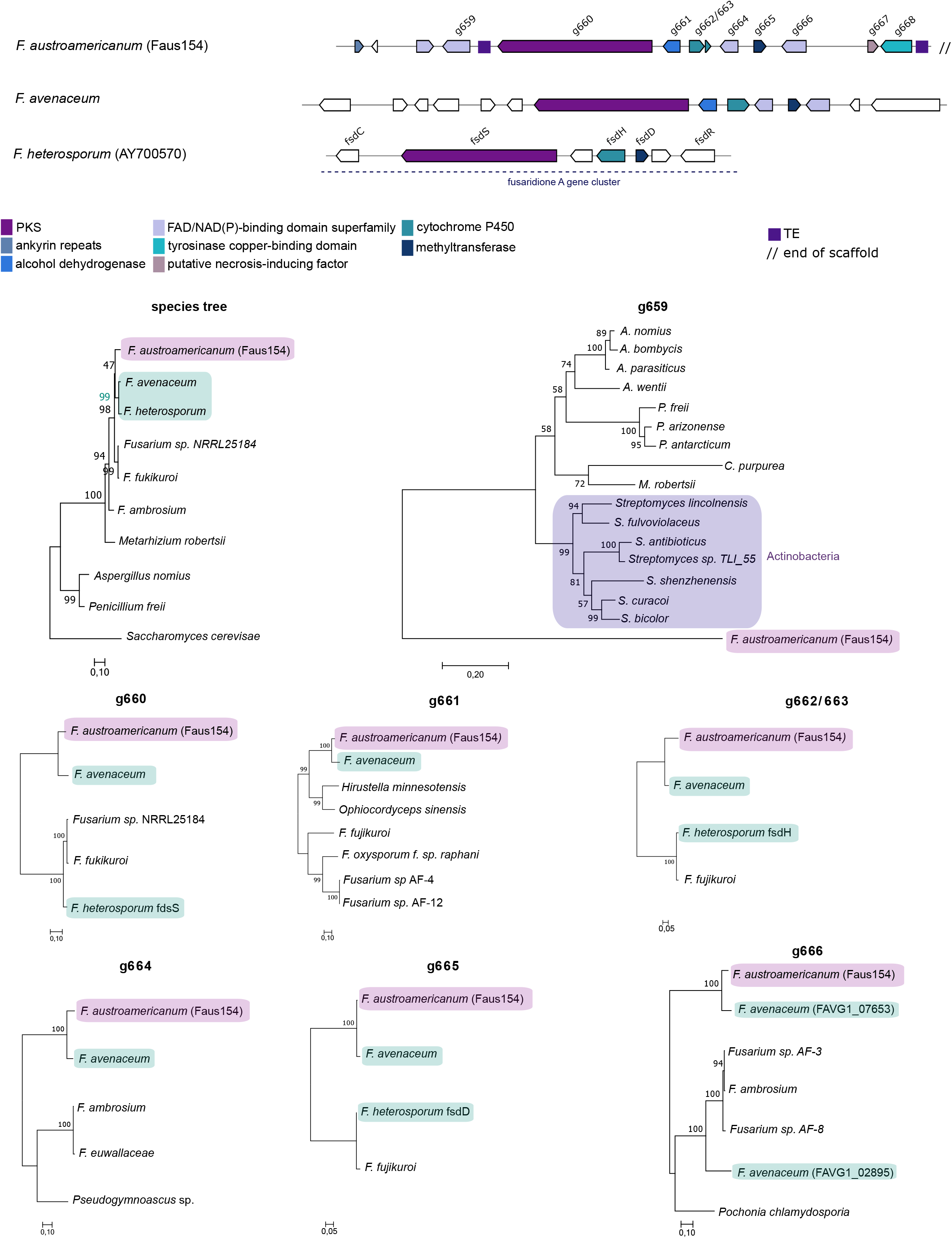
Synteny plot of the SM54 gene cluster. Arrows of identical color correspond to homologous genes and identify the predicted protein function. White arrows identify genes without a homolog in corresponding strain. Phylogenetic trees were built using maximum likelihood and the JTT matrix-based model with 1000 boostrap replicates. The species tree was based on the concatenated genes EF-1α, RPB1 and RPB2. *Saccharomyces cerevisiae* was used as the outgroup.

Interestingly, a second gene of the SM54 cluster (Faus154_g659), encoding a NAD(P)/FAD-binding protein was gained most likely through horizontal transfer from bacteria. A homology search identified a homolog in the Actinobacteria *Streptomyces antibioticus* with 44.3% identity and 56.8% similarity followed by several other *Streptomyces* spp. strains as the next best hits (Supplementary Table S4). The homologs in *F. austroamericanum* and *S. antibioticus* share the same NAD(P)/FAD-binding domains (Supplementary Figure S2). Among fungi, hits to the *F. austroamericanum* homolog were of lower percentage identity, the best hit was found in the ascomycete *Aspergillus wentii* with 40.6% identity (Supplementary Table S4). Hence, this suggests a more recent horizontal transfer event between an ancestor of *Streptomyces* and *Aspergillus.* The lack of close orthologues of Faus154_g659 in other fungi of the same class (Sordariomycetes) and the amino acid and functional homology found in bacteria, suggested an ancient bacterial origin of this gene via a horizontal transfer event.

### Gene cluster reconfiguration across diverse fungi

The cluster SM53 is shared among two FGSC strains, *F. cortaderiae* (strain Fcor153) and *F. austroamericanum* (strain Faus151). In the second *F. austroamericanum* strain (Faus154), the cluster is missing most genes and suffered pseudogenization (Figure 6). We conducted a broad homology search across fungi and found SM53 to be present in *F. bulbicola*, which is not a member of the FGSC. In *F. bulbicola*, the core gene set clusters with at least six additional genes that are typically associated with a fumonisin gene cluster including a cytochrome P450 homologue identified as the fumonisin gene *cpm1*. Even though *F. bulbicola* is a fumonisin C producer, the specific strain was identified as a non-producer (Brown and Proctor 2016). To investigate possible gaps in the genome assembly near the gene cluster, we searched the *F. bulbicola* genome for additional fumonisin genes. We analyzed homology at the nucleotide and amino acid level between *F. bulbicola* and the *F. oxysporum* strain RFC O-1890. RFC O-1890 is a fumonisin C producer and the most closely related available strain to *F. bulbicola* (Supplementary Table S5) (Proctor et al. 2008). We identified fumonisin cluster elements on 4 different *F. bulbicola* scaffolds with the exception of *FUM11* and *FUM17*.

**Figure 6.**
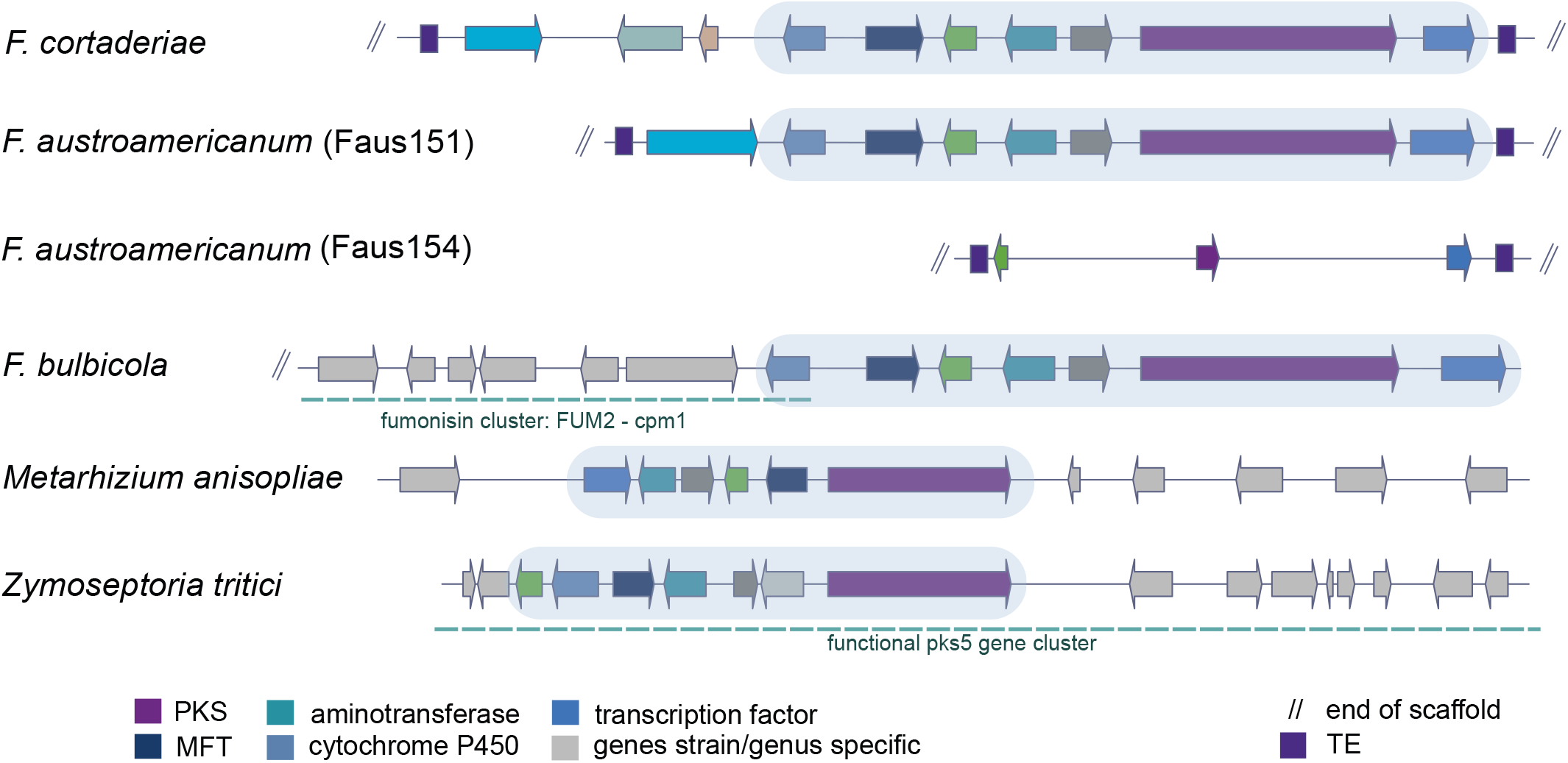
Synteny plot of the SM53 gene cluster. Arrows of identical color correspond to homologous genes and identify the predicted protein function. Light gray arrows correspond to genes lacking homology among analyzed strains. Light blue identifies the conserved core set of genes. Blue dotted lines in *Fusarium bulbicola* correspond to the fumonisin cluster adjacent to the core set and in *Zymoseptoria tritici* to the PKS5 gene cluster upregulated during infection in wheat. PKS: polyketide synthase; MFT: major facilitator superfamily transporter; TE: transposable element.

We found additional evidence for the SM53 core cluster in distantly related fungi including *Metarhizium, Aspergillus* and *Zymoseptoria*. The cluster variant identified in the entomopathogenic fungus *M. anisopliae* was identified as a Mapks12 cluster (Sbaraini, et al. 2016). Although, the full cluster size in *M. anisopliae* is still unknown, transcriptomic data showed expression of the gene encoding the PKS and adjacent genes in culture media (Sbaraini et al. 2016). In the wheat pathogen *Z. tritici*, the core gene set is forming a larger functional cluster and transcriptomic data shows coordinated upregulation, and high expression upon infection of wheat (Palma-Guerrero et al. 2016). Phylogenetic analyses of the backbone gene encoding a PKS showed broad congruence with the species tree consisted with long-term maintenance despite widespread losses in other species (Supplementary Figure S3). The highly conserved core cluster segment may constitute a functional cluster because it encodes a typical complement of cluster functions including a PKS, a cytochrome P450, a dehydrogenase, a methyltransferase, a transcription factor and a major facilitator superfamily transporter.

### Transposable elements associated with gene cluster rearrangements

We found evidence for the gene cluster SM48 in four different species of the FGSC (*F. cortaderiae, F. austroamericanum, F. meridionale* and *F. asiaticum*). In *F. graminearum* s.s., the PKS backbone gene is absent. However, we found evidence for five additional genes of SM48 in four different chromosomal locations and two different chromosomes (Figure 7). A gene encoding a homeobox-like domain protein, a transporter gene and the flanking genes clustered together on chromosome 2, but in two different loci at approximately 60 kb and 50 kb from each other, respectively. The gene encoding the glycosyl hydrolase, which is next to the backbone gene encoding the PKS in the canonical SM48 gene cluster configuration, was found as an individual gene in the subtelomeric region of chromosome 4. *F. avenaceum* is the only analyzed species outside the FGSC that shared the PKS gene (Figure 7). Interestingly, the SM48 gene cluster contained a series of transposable elements integrated either next to the gene encoding the PKS and/or the gene encoding the glycosyl hydrolase. Furthermore, a phylogenetic analysis showed a patchy taxonomic distribution of homologues across the *Fusarium* genus (Supplementary Table S6). The gene cluster SM48 was most likely vertically inherited by the FGSC because both *F. avenaceum* and *F. culmorum* showed rearranged configurations compared to FGSC species. Disrupted cluster variants are present in the clade formed by *F. graminearum* s.s, *F. boothi*, *F. louisianense* and *F. gerlachii*. The high density of transposable elements might have facilitated the rearrangement of the gene cluster.

**Figure 7.**
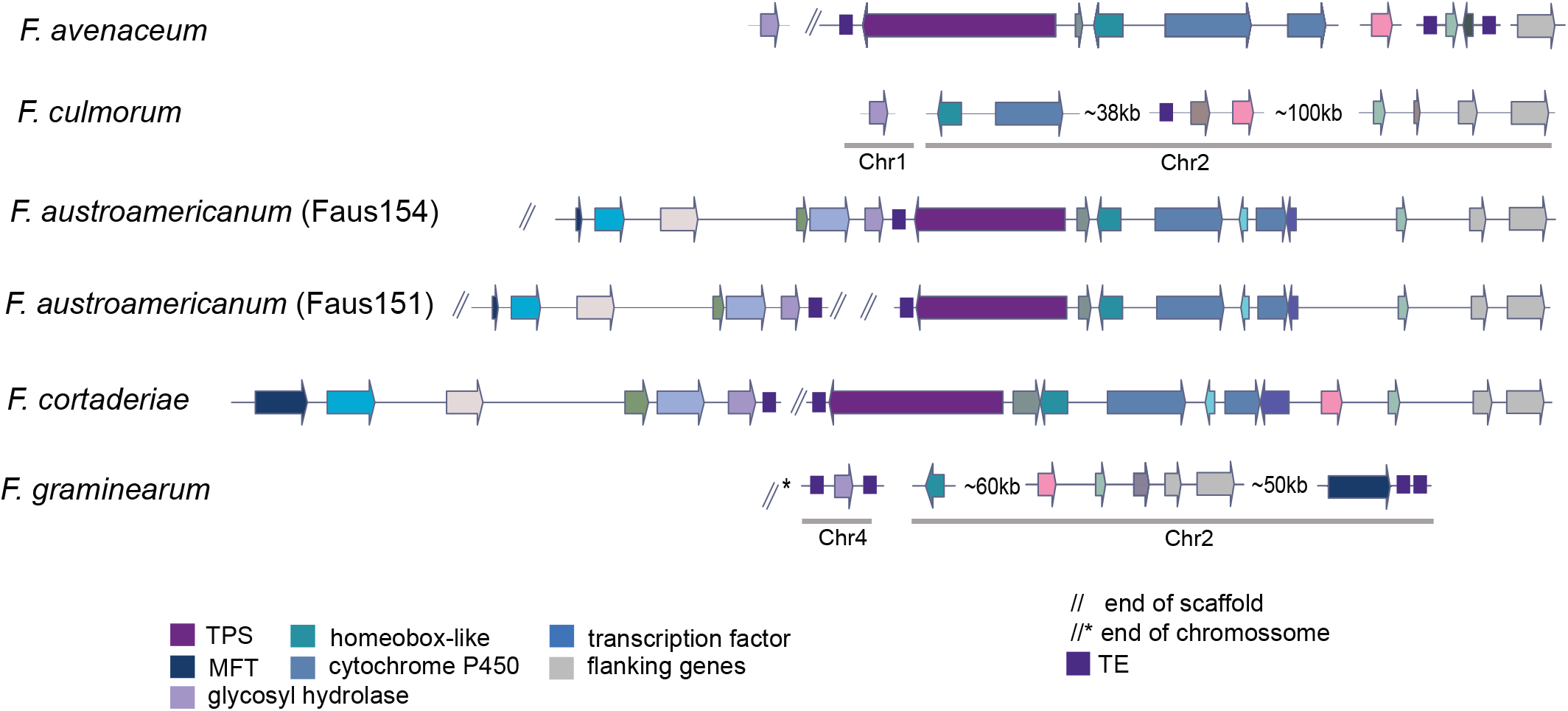
Synteny plot of the SM48 gene cluster. Arrows of identical color correspond to homologous genes and identify the predicted protein function. Values adjacent to disrupted clusters define physical distances and grey bars below genes define to chromosomal locations. TPS: terpene synthase; MFT: major facilitator superfamily transporter; TE: transposable element.

### Transposable elements families in FGSC

Several gene clusters of category 2 and 3 (SM46, SM48, SM48 and SM54; Figure 2), which showed various levels of reconfigurations were flanked by transposable elements. To understand broadly how transposable elements may have contributed to gene cluster evolution, we analyzed the identity of transposable elements across the genomes and in close association with gene clusters. We found overall no difference in transposable element density in proximity to gene clusters compared to the rest of the genome with the exception of the *F. asiaticum* strain FasiR (Supplementary Figure S4). FasiR showed about twice the transposable element density in proximity to clusters (9.9%) compared to genome-wide average (4.1%). Next, we analyzed the frequency of individual transposable element families within 10 kb of gene clusters and compared this to the frequency in all 10 kb windows across the genomes of the FGSC (Figure 8A). We found a series of transposable element families that were more frequent in proximity to gene clusters (Figure 8B). The most abundant elements in the genomes of the FGSC are the unclassified elements 3-family-62 (mean frequency of 0.147 per 10 kb window) followed by 2-family-17 (mean frequency of 0.124). In proximity to SM gene clusters, the frequency of the 2-family-17 was higher than 3-family-62 in 54% of the strains, with an overall mean of 0.174 and 0.160, respectively. The element 4-family-882, which is enriched in the clade comprising *F. graminearum* s.s*, F. gerlachii, F. boothi* and *F. louisianense*, as well as the strain *F. cortaderiae*, is seven times more frequent near SM gene clusters compared to the whole genome (FgramR; Figure 8B).

**Figure 8.**
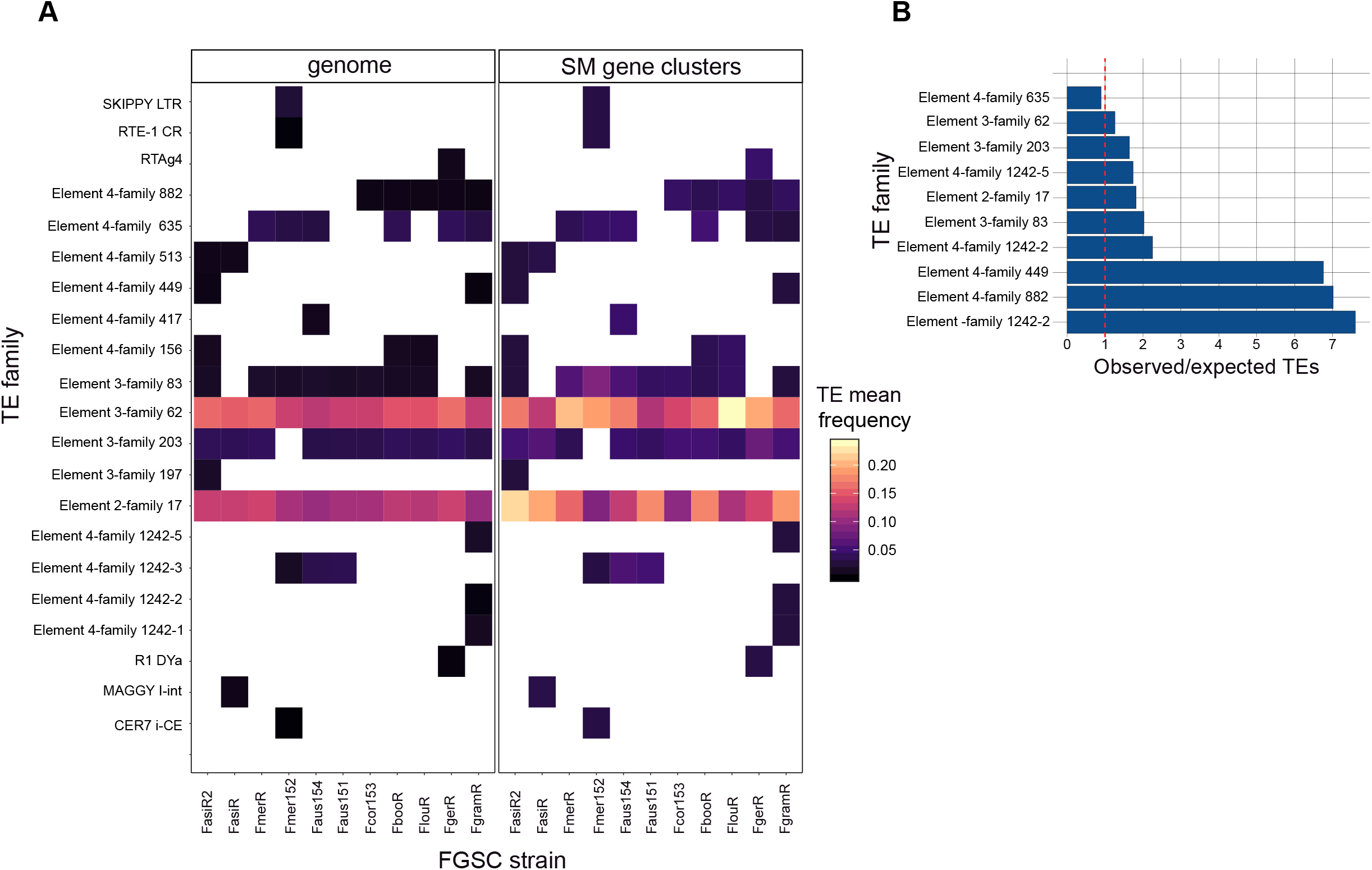
A - Heatmap with the most frequent transposable element families flanking FGSC gene clusters and the overall genome in 10 kb windows. B-Bar chart showing the ratio of the observed (SM gene cluster) over the expected transposable elements (genome) in the *F. graminearum* reference genome (FgramR). Red dotted line marks the ratio of one representing no difference.

## Discussion

We assembled and analyzed a comprehensive set of genomes representative of the FGSC diversity. Our phylogenomic analyses corroborated previous multilocus studies and refined our understanding of the evolutionary relationships within the complex (O’Donnel et al, 2004; Aoki et al. 2012). The recent speciation among members of the FGSC led to differentiation in host range, genome size, gene and transposable element content. Our analyses of SM gene clusters within the FGSC revealed more complexity than previously reported (Walkowiak et al. 2016). Individual gene clusters underwent independent gene losses, sequence rearrangements associated with transposable elements and multiple horizontal transfer events, leading to presence/absence polymorphism and chemical diversity within the FGSC.

### A diverse SM gene cluster pangenome of the FGSC

We performed pangenome analyses of eight species of FGSC (11 isolates) to exhaustively characterize the presence of known and unknown SM gene clusters. The emergence of the FGSC was accompanied by the loss and rearrangement of several SM gene clusters. The most recent common ancestor with other members of the *Fusarium* clade likely carried more SM gene clusters. The recently lost clusters may underlie the adaptation to wheat as a primary host. Among the fully conserved gene clusters within the FGSC, we found clusters underlying the production of siderophores including triacetylfusarin and ferricrocin that facilitate iron acquisition (Charlang et al. 1981). We also found conserved clusters underlying the production of virulence factors, *e.g.* gramillin on maize (Bahadoor et al. 2018). The conservation likely reflects the essential functions of these metabolites in the life cycle of the fungi. The SM gene clusters not fixed within the FGSC spanned a surprisingly broad number of types including TPS, NRPS, NRPS-TPS, and NRPS-PKS. Segregating gene clusters may reflect adaptation to niches specific to a subset of the FGSC. Such adaptation may explain the conservation of the apicidin cluster in the *F. asiaticum* strain FasiR2 isolated from maize and the lack of the cluster in the strain FasiR isolated from barley (O’Donnel et al. 2000).

How the environmental heterogeneity selects for diversity in SM gene clusters among closely related species is poorly understood, yet studies have found strong associations of SM gene clusters with different lifestyles and geographical distribution (Reynolds, et al. 2017, Wollenberg et al. 2018). The fusaristatin A gene cluster, thought to be missing in *F. pseudograminearum* (but present in FGSC), was recently found to be functional in a Western Australian population of *F. pseudograminearum* (Wollenberg et al. 2018). In FGSC, trichothecenes are key adaptations to exploit the host. Different forms of trichothecenes (*i.e.* deoxynivalenol, 3-acetyldeoxynivalenol, 15-acetyldeoxynivalenol and nivalenol chemotypes) are segregating in pathogen populations due to balancing selection (Ward et al. 2002). The trichothecene polymorphism is likely adaptive with the role in pathogenesis depending both on the crop host (Desjardins et al. 1992; Proctor et al. 2002; Cuzick et al. 2008) and the specific trichothecene produced (Carter et al. 2002, Ponts et al. 2009; Spolti et al. 2012). For example, nivalenol production is associated with pathogenicity on maize and deoxynivalenol is essential to Fusarium head blight in wheat spikelets but seems to play no role for pathogenicity on maize (Maier et al. 2006). Both toxins play no role in pathogenicity on barley. A variable pangenome of metabolic capacity maintained among members of the FGSC may, hence, also serve as a reservoir for adaptive introgression among species.

### Mechanisms generating chemical diversity in *Fusarium*

Our study revealed a complex set of mechanisms underlying SM gene cluster diversity in FGSC. We found that multiple independent losses are a key factor generating extant cluster diversity within the FGSC and *Fusarium*. The SM43 (guaia,6-10(14)-diene) and the apicidin clusters were lost multiple times within *Fusarium* and in different lineages of the FGSC. Independent losses are frequently associated with the evolutionary trajectory of SM gene clusters (Patron et al. 2007; Khaldi et al. 2008). The evolution of the galactose (GAL) cluster in yeasts was characterized by multiple independent losses and at least 11 times among the subphyla of *Saccharomycotina* and *Taphrinomycotina* (Riley at al. 2016). Similarly, Campbell et al. (2012) showed that the bikaverin gene cluster was repeatedly lost in the genus *Botrytis* after receiving the cluster horizontally from a putative *Fusarium* donor. A gene cluster loss is typically favored by either a decreased benefit to produce the metabolite or an increase in production costs (Rokas et al. 2018). Along these lines, the *black queen hypothesis* conveys the idea that the loss of a costly gene (cluster) can provide a selective advantage by conserving an organism’s limited resources (Morris et al. 2012). Such loss-of-function mutations (e.g abolishing metabolite production) are viable in an environment where other organisms ensure the same function (Mas et al. 2016; Morris et al. 2012). The *black queen hypothesis* may at least partially explain the metabolite diversity and high level of cluster loss in the FGSC if different lineages and species frequently co-exist in the same environment or host.

Horizontal gene transfer is an important source of gene cluster gain in fungi (Kaldhi et al. 2008; Khaldi and Wolfe, 2011; Slot and Rokas, 2011; Campbell et al. 2012; Slot and Rokas, 2012) and likely contributed to the FGSC gene cluster diversity. Here, we report an unusual case of multiple, independent horizontal transfer events involving an ancient transfer from bacteria and a more recent fungal donor. The horizontal transfer contributed to the formation of the SM54 gene cluster found in the strain *F. austroamericanum* (Faus154). Horizontal transfer events have been proposed as an important form of pathogenicity emergence. A gene cluster of *F. pseudograminearum* was most likely formed by three horizontally acquired genes from other pathogenic fungi. An additional gene of the cluster encoding an amidohydrolase was received from a plant-associated bacterial donor and associated with pathogenicity on wheat and barley (Gardiner et al. 2012). Similarly, the *Metarhizum* genus of entomopathogens acquired at least 18 genes by independent horizontal transfer events that contribute to insect cuticle degradation (Zhang et al. 2018).

Our analyses revealed the SM53 gene cluster core segment that is conserved across distantly related genera. The core section underlies the formation of superclusters through the rearrangement with a separate cluster and likely led to neofunctionalization. The backbone and adjacent genes in the conserved segment were found to be expressed in *M. anisopliae* in culture medium (Sbaraini et al. 2016). In the wheat pathogen *Z. tritici*, the core segment was associated with additional genes forming a larger cluster with coordinated upregulation upon host infection (Palma-Guerrero et al. 2016). A study in *A. fumigatus* identified a similar event, where the clusters underlying pseurotin and fumagillin production were rearranged to form a supercluster (Wiemann et al. 2013). Similar to the gene cluster SM53, the segments of the supercluster were conserved in *A. fischeri* and in the more distantly related species *M. robertsii*. Taxonomically widespread conserved gene cluster segments may represent functional but transitory gene cluster variants that can give rise to superclusters. Viable, transitory stages are an efficient route to evolve new metabolic capacity across fungi (Rokas et al. 2018, Lind et al. 2017).

### Transposable elements as drivers of gene cluster rearrangements

Our analyses revealed that gene cluster gains and losses in the FGSC were associated to transposable elements. We found an enrichment in transposable elements adjacent or integrated within different clusters (*i.e.* SM1, SM21, SM48, SM53 and SM54). Our data strongly suggests that the cluster SM48 emerged within FGSC and suffered transposable element-associated chromosomal rearrangements in the *F. graminearum* s.s clade followed by functional loss. The SM53 pseudogenization and gene loss in the *F. austroamericanum* strain Faus154 was likewise caused by transposable elements insertions adjacent to the cluster. Transposable elements play an important role in the evolution, particularly related to virulence, of fungal pathogens (Gardiner et al. 2013; Sánchez-Vallet et al. 2018; Fouché et al. 2018). Transposable elements can induce gene cluster rearrangements due to non-homologous recombination among repeat copies (Boutanaev and Osbourne 2018), but also impact genome structure and function by causing gene inactivation, copy number variation, and expression polymorphism (Manning et al. 2013; Sánchez-Vallet et al. 2017; Krishnan et al. 2018). For example, flanking transposable elements likely caused transposition events of a specialized cluster in *A. fumigatus* (Lind et al. 2017). The enriched transposable elements near gene clusters in FGSC genomes were likely overall an important driver of gene cluster loss, rearrangement, and neofunctionalization.

Our study provides insights into the evolutionary origins of SM gene clusters in a complex of closely related species. The recency of speciation within the FGSC is reflected by the predominant number of conserved gene clusters. Nevertheless, the FGSC accumulated previously under-appreciated gene cluster diversity, which originated from a broad spectrum of mechanisms including parallel gene losses, rearrangements and horizontal acquisition. Independent losses within the complex were likely due to ecological drivers and strong selection. Hence, environmental heterogeneity may play an important role in gene cluster evolution (Rokas et al. 2018). Chromosomal rearrangements underlying cluster loss were often complex and were likely facilitated by transposable elements. At the same time, chromosomal rearrangements contributed to gene cluster neofunctionalization. The extant chemical diversity of FGSC highlights the importance of transitory stages in the evolution of specialized metabolism among very closely related species.

## Author contributions

SMT, LOR, BC and DC conceived the study; SMT, LOR and BC provided samples and datasets; SMT, UO analyzed the data; SMT and DC wrote the manuscript; LOR, UO and BC edited the manuscript

## Data availability

All raw sequence data was uploaded to the NCBI Short Read Archive (PRJNA542165).

## Acknowledgments

We thank Dr. Robert Proctor from the National Center for Agricultural Utilization Research (United States Department of Agriculture) for kindly providing the genomic sequences of *F. bulbicola.* This research was supported by FAPESP (Fundação de Amparo a Pesquisa do Estado de São Paulo) grant process 2017/22369-7 and 2016/04364-5. DC receives support from the Swiss National Science Foundation (grants 31003A_173265 and IZCOZO_177052).

