## Supplementary Figure for "Complex evolutionary origins of specialized metabolite gene cluster diversity among the plant pathogenic fungi of the *Fusarium graminearum* species complex"

### Supplementary Figures

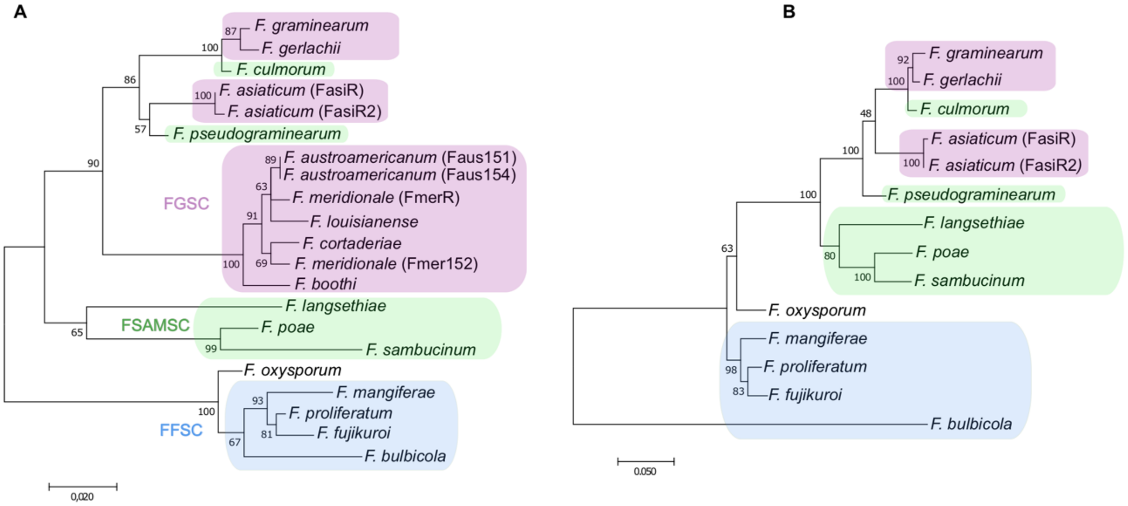


**Supplementary Figure S1.** Phylogenetic analysis of the genes composing the SM43 gene cluster underlying guaia-6,10(14)-diene metabolite production. A: pyoverdine biosynthesis gene tree. B: TPS gene tree. The trees were built using maximum likelihood and the JTT matrix-based amino acid model with 1000 bootstrap replicates. FGSC: *Fusarium graminearum* species complex; FSAMSC: *Fusarium sambucinum* species complex; FFSC: *Fusarium fujikuroi* species complex. The species tree is shown in Figure 1.

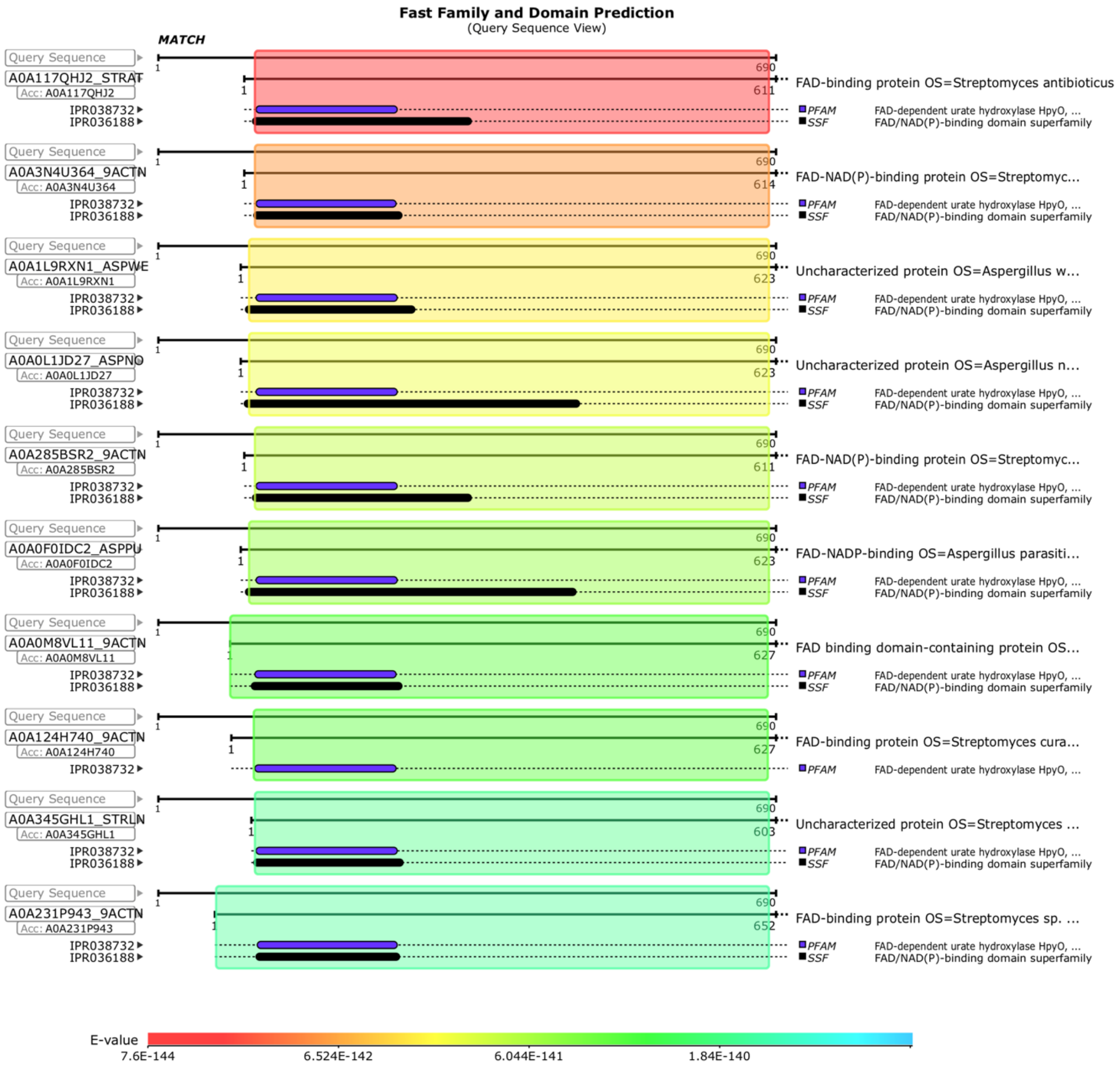


**Supplementary Figure S2.** Functional prediction of the gene Faus_gene659 based on the EMBL-EBI BLAST protein similarity search engine. Results are ordered according to the best e-value score.

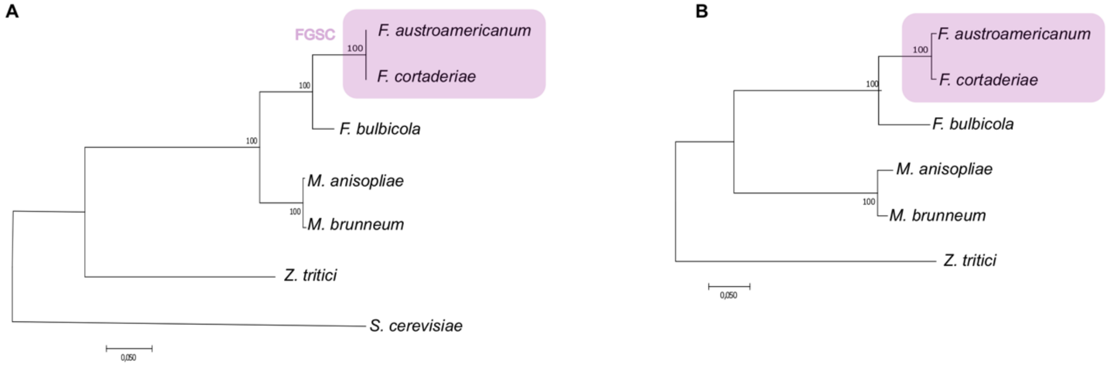


**Supplementary Figure S3.** Phylogenetic analyses of the genes composing the SM48 gene cluster. A: species tree based on the concatenated genes EF-1α, RPB1 and RPB2. *Saccharomyces cerevisiae* was used as the outgroup. B: PKS gene tree. The trees were built using maximum likelihood and the JTT matrix-based amino acid model with 1000 bootstrap replicates. FGSC: *Fusarium graminearum* species complex.

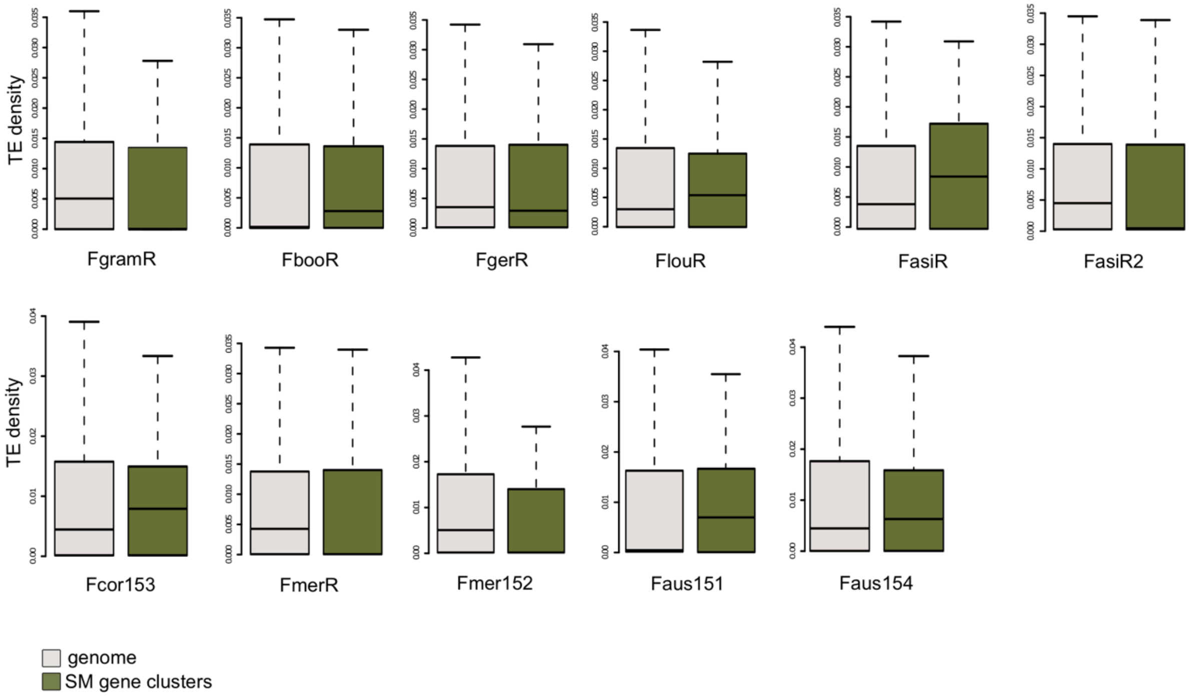


**Supplementary Figure S4.** Comparison of the genome-wide transposable element (TE) density in 10 kb windows and flanking regions (10 kb) of the secondary metabolite (SM) gene clusters. The black line in the boxplot corresponds to the median.

### Supplementary Table legends

[see separate file]

**Supplementary Table S1.** Overview of the analyzed genomes. The colored section corresponds to the members of the *Fusarium graminearum* species complex.

**Supplementary Table S2.** Strain accession numbers.

**Supplementary Table S3.** List of the specialized metabolite (SM) gene cluster pangenome of the *Fusarium graminearum* species complex. * SM gene clusters were considered conserved when the backbone gene showed a homology above 70% and was present in at least 24 out of 26 analyzed *Fusarium* genomes.

**Supplementary Table S4.** Results of the blastp top hits for the SM54 gene cluster genes.

**Supplementary Table S5.** Percent identity of the fumonisin (FUM) gene cluster orthologues in *Fusarium oxysporum* and *F. bulbicola*. Gene homology was determined using blastn and blastp analyses between the *F. oxysporum* strain FRC O-1890 (accession: EU449979.1) and *F. bulbicola* (FbulR). [a] Values were determined using the nucleotide sequences of the coding regions (CDS) and introns. [b] no hit found on the scaffold in blastp analyses.

**Supplementary Table S6.** Gene homology of the SM48 cluster genes based on blastp among the analyzed *Fusarium* species. The colored section corresponds to different scaffolds or chromosomes in the genome. na – not available. Homology was determined based on identity above 70% and synteny conservation among neighboring genes.
